# A max-margin training of RNA secondary structure prediction integrated with the thermodynamic model

**DOI:** 10.1101/205047

**Authors:** Manato Akiyama, Kengo Sato, Yasubumi Sakakibara

## Abstract

**Motivation:** A popular approach for predicting RNA secondary structure is the thermodynamic nearest neighbor model that finds a thermodynamically most stable secondary structure with the minimum free energy (MFE). For further improvement, an alternative approach that is based on machine learning techniques has been developed. The machine learning based approach can employ a fine-grained model that includes much richer feature representations with the ability to fit the training data. Although a machine learning based fine-grained model achieved extremely high performance in prediction accuracy, a possibility of the risk of overfitting for such model has been reported.

**Results:** In this paper, we propose a novel algorithm for RNA secondary structure prediction that integrates the thermodynamic approach and the machine learning based weighted approach. Our fine-grained model combines the experimentally determined thermodynamic parameters with a large number of scoring parameters for detailed contexts of features that are trained by the structured support vector machine (SSVM) with the *ℓ*_1_ regularization to avoid overfitting. Our benchmark shows that our algorithm achieves the best prediction accuracy compared with existing methods, and heavy overfitting cannot be observed.

**Availability:** The implementation of our algorithm is available at https://github.com/keio-bioinformatics/mxfold.

## 1 Introduction

Non-coding RNAs (ncRNAs) that are not translated into proteins were formerly considered as junk regions. However, these various functions have been revealed in recent years ranging from the process of development and cell differentiation to the cause of disease. Since the functions of ncRNAs are believed to be closely related to the structures of ncRNAs, it is possible to infer their biological functions from their structures. RNA tertiary structures can be determined by experimental assays including X-ray crystal structure analysis and nuclear magnetic resonance (NMR). However, there are severe difficulties of these experimental assays such as high experimental cost and low throughput. In addition, the computational techniques to predict RNA tertiary structures have still been immature. Therefore, the computational prediction of RNA secondary structures, which can be easily modeled by a set of hydrogen bonds between nucleotides, has frequently been used instead.

From the viewpoint of the scoring scheme, RNA secondary structure prediction methods are roughly classified into three approaches: a thermodynamic approach, a probabilistic approach, and a weighted approach Rivas (2013). The thermodynamic approach has been the most popular approach that finds a thermodynamically most stable secondary structure with the minimum free energy (MFE) and has been utilized by a number of tools including UNAfold Zuker (1989), RNAfold Lorenz *et al.* (2011), and RNAstructure Reuter and Mathews (2010). RNA secondary structures can be decomposed into characteristic substructures such as hairpin loops and base-pair stacking according to the nearest neighbor model Zuker and Stiegler (1981). Free energy of each substructure was determined by experimental methods such as the optical melting experiment Schroeder and Turner (2009). The free energy of the secondary structure is calculated by summing up the free energy of each substructure in the secondary structure. The dynamic programming technique enables us to efficiently find the MFE structure from all possible secondary structures for a given RNA sequence.

The probabilistic approach has employed generative models including stochastic context-free grammars (SCFGs) for modeling RNA secondary structures. SCFGs are defined by a set of derivation rules, or grammar, whose probabilities are trained by the maximum likelihood (ML) estimation from the training data, and were applied to RNA secondary structure prediction Sakakibara *et al.* (1994); Eddy and Durbin (1994); Knudsen and Hein (1999); Dowell and Eddy (2004). Sato *et al.* proposed a non-parametric Bayesian extension of SCFGs with the hierarchical Dirichlet process that can find an optimal RNA grammar from the training data Sato *et al.* (2010). Rivas *et al.* developed a framework called TORNADO for flexibly describing RNA grammars, and showed that a complex RNA grammar that simulates the nearest neighbor model can achieve as accurate predictions as the weighted models can Rivas *et al.* (2012).

The weighted approach has utilized machine learning techniques instead of the experimental techniques in order to determine weights for decomposed substructures, i.e., scoring parameters. CONTRAfold was developed based on the conditional log-linear models (CLLMs) that find scoring parameters that can most probably discriminate between correct structures and incorrect structures Do *et al.* (2006). Simfold implemented Boltzmann likelihood algorithm with feature relationships between parameters (BL-FR), which is similar to CLLMs, but incorporated free energy parameters Andronescu *et al.* (2010). ContextFold employed a fine-grained model that includes much richer contexts of features with the ability to fit the training data, combined with a machine learning algorithm Zakov *et al.* (2011). Although ContextFold achieved extremely high performance in prediction accuracy, Rivas *et al.* reported a possibility of the risk of overfitting for ContextFold Rivas (2013). From this observation, we can see that an important issue for further improving prediction accuracy is to effectively learn a large number of scoring parameters for a fine-grained model without overfitting.

In this paper, we propose a novel algorithm for RNA secondary structure prediction that integrates the thermodynamic approach and the machine learning based weighted approach. Our fine-grained model combines the experimentally-determined thermodynamic parameters with a large number of scoring parameters for detailed contexts of features. In order to train the scoring parameters of the fine-grained model, we employ the structured support vector machine (SSVM) Tsochantaridis *et al.* (2005) with the *l*_1_ regularization to avoid overfitting. Our benchmark shows that our algorithm achieves the best prediction accuracy compared with existing methods, and heavy overfitting as shown in ContextFold cannot be observed.

The major advantages of our work are summarized as follows: (i) The max-margin based training algorithm learns our fine-grained model that can perform accurate secondary structure prediction, and (ii) our scoring model that integrates the thermodynamic and machine learning based model enables accurate and robust structure prediction even for unobserved substructures in the training dataset.

## 2 Methods

### 2.1 Preliminaries

Let Σ = {A, C, G, U} and Σ* denote the set of all finite RNA sequences consisting of bases in Σ. For a sequence *x* = *x*_1_*x*_2_ … *x*_*n*_ ∈ Σ*, let |*x*| denote the number of symbols appearing in *x*, which is called the length of *x*. Let *S*(*x*) be a set of all possible secondary structures of *x*. A secondary structure *y* ∈ *S*(*x*) is represented as a |*x*| × |*x*| binary-valued triangular matrix *y* = (*y*_*ij*_)_*i*<*j*_, where *y*_*ij*_ = 1 if and only if bases *x*_*i*_ and *x*_*j*_ form a base-pair composed by hydrogen bonds including the Watson-Crick base-pairs (A-U and G-C), the Wobble base-pairs (G-U).

### 2.2 Scoring model

A scoring model *f*(*x*, *y*) is a function that assigns real-valued scores to an RNA secondary structure *y* ∈ *S*(*x*) for an RNA sequence *x* ∈ Σ*. Our aim is to find a secondary structure *y* ∈ *S*(*x*) that maximizes the scoring function *f*(*x*, *y*) for a given RNA sequence *x* ∈ Σ*.

RNA secondary structures can be decomposed into characteristic substructures, or features, such as hairpin loops and base-pair stacking. We denote by Φ(*x*, *y*) the feature representation vector of (*x*, *y*), which consists of the number of occurrence of every feature in (*x*, *y*). Each feature in Φ is associated with a corresponding score or weight. We assume a linear scoring model of RNA secondary structures as:

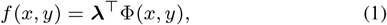

where λ is a weight vector in which λ_*i*_ is the weight of the *i*-th feature in Φ.

Note that the thermodynamic approach can be represented by this linear scoring model if we define Φ as the nearest neighbor model and the corresponding weights as the negative of experimentally determined free energy parameters.

We propose a novel scoring model that integrates the thermodynamic approach and the machine learning based weighted approach. We define our scoring model as:

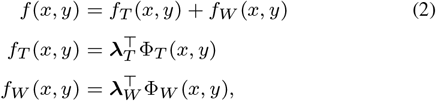

where *f*_*T*_(*x*, *y*) (resp. *f_W_* (*x*, *y*)) is the contribution of the thermodynamic model (resp. the machine learning model) to our scoring model. For the thermodynamic model, we employ the nearest neighbor model as Φ_*T*_ and the negative of the Turner free energy parameters Turner and Mathews (2010) as λ_*T*_. For the machine learning model, we construct a fine-grained model as Φ_*W*_ (see Sec. 2.3) and corresponding weights λ_*W*_ that are trainable from training data by using SSVM (see Sec. 2.5).

### 2.3 Feature representations

Both feature representations Φ_*T*_ and Φ_*W*_ are based on the nearest neighbor model Zuker and Stiegler (1981), including base helices, dangling ends, terminal mismatches, hairpin loops, bulge loops, internal loops, multibranch loops and external loops (Fig. 1).

**Fig. 1.**
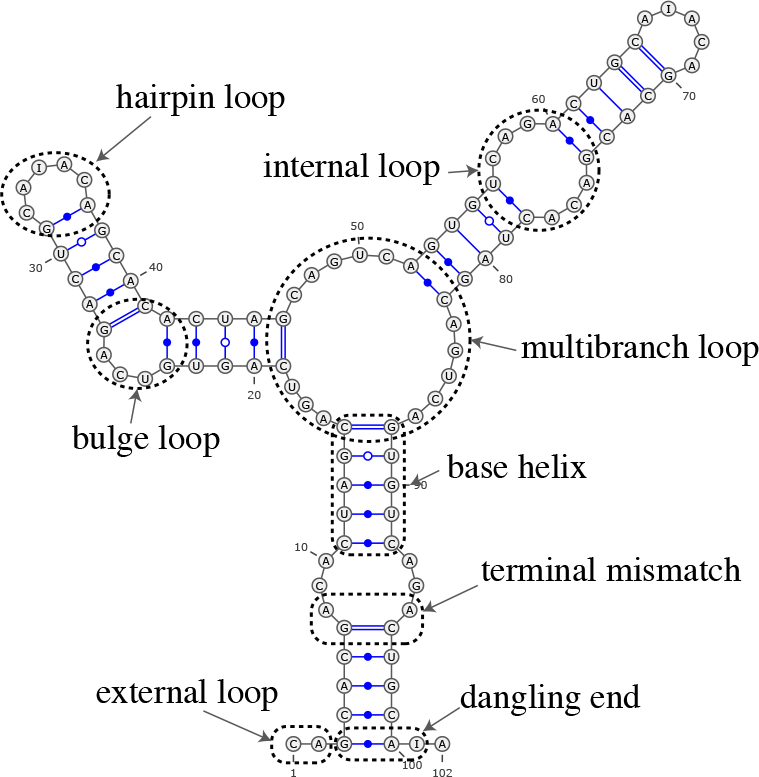
Examples of substructures defined in the standard nearest neighbor model.

In order to calculate the free energy of RNA secondary structures more precisely, some specialized loop parameters have been adopted in frequently used free energy parameter sets for the standard nearest neighbor model. For example, the Turner 1999 and 2004 models contain several sequential features such as hairpin loops with 3, 4 or 6 nucleotides and internal loops with (1, 1) nucleotides (1 nucleotide at 5’ loop and 1 nucleotide at 3’ loop), (1, 2) nucleotides and (2, 2) nucleotides Turner and Mathews (2010).

As the fine-grained feature representation Φ_*W*_, we employ much longer sequential features for hairpin loops with *m* nucleotides, bulge loops with m nucleotides and internal loops with (*m*, *n*) nucleotides (*m* ≤ *L* and *m* + *n* ≤ *L*) in addition to the standard nearest neighbor model. We use *L* = 7 by default as described in Results. See Sec. 3.5 for more details.

### 2.4 Decoding algorithm

#### 2.4.1 Viterbi decoding

Since both Φ_*T*_ and Φ_*W*_ are based on the nearest neighbor model, any secondary structures can be decomposed into the same substructures for both representations. Therefore, we can find the most probable secondary structure that maximizes Eq. (2) by the Zuker-style dynamic programming algorithm Zuker and Stiegler (1981).

#### 2.4.2 Posterior decoding

The posterior probability of the secondary structure *y* given RNA sequence *x*, *p*(*y* | *x*), under the scoring model *f*(*x*, *y*) is calculated by:

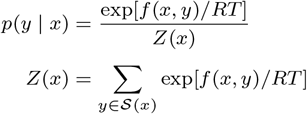

where *R* is the gas constant and *T* is the absolute temperature. The base-pairing probability *p*_*ij*_ is the probability that the *i*-th and *j*-th nucleotides form a base-pair, which is defined as follows:

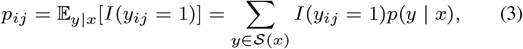

where *I*(*condition*) is an indicator function which takes a value of 1 or 0 depending on whether the *condition* is true or false. The McCaskill algorithm McCaskill (1990) can be utilized to efficiently calculate the base-pairing probabilities (3) by the dynamic programming techniques.

We define a gain function between a true structure *y* and a candidate structure *ŷ* by

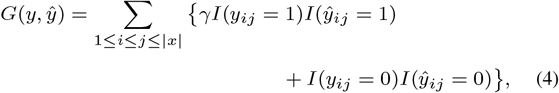

where *γ* > 0 is a weight for base-pairs. The gain function (4) is equal to the weighted sum of the number of true positives and the number of true negatives of base-pairs.

The expectation of the gain function (4) with respect to an ensemble of all possible secondary structures under a given posterior distribution *p*(*y* | *x*) is

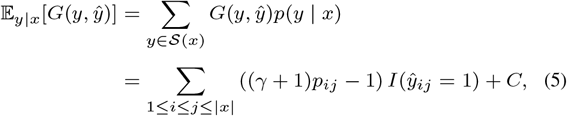

where *C* is a constant independent of *ŷ*.

Then, we can find *ŷ* that maximizes the expected gain (5) using the recursive equations:

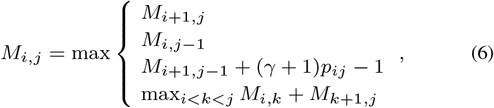

and tracing back from *M*_1, |*x*|_.

We can control the trade-off between specificity and sensitivity by *γ*. We call the maximization of Eq. (5) the generalized centroid estimator (GCE) since this is equivalent to the centroid estimator Ding *et al.* (2005); Carvalho and Lawrence (2008) for *γ* = 1. The generalized centroid estimator is very similar to the maximum expected accuracy (MEA) estimator Do *et al.* (2006). See Hamada *et al.* (2009); Sato *et al.* (2009) for more details.

### 2.5 Learning algorithm

To optimize the feature parameter λ_*W*_, we employ a max-margin framework called structured support vector machines (SSVM) Tsochantaridis *et al.* (2005). Given a training dataset 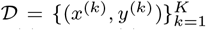, where *x*^(*k*)^ is the *k*-th RNA sequence and *y*^(*k*)^ ∈ *S*(*x*^(*k*)^) is the correct secondary structure for the *k*-th sequence *x*^(*k*)^, we aim to find λ_*W*_ that minimizes the objective function

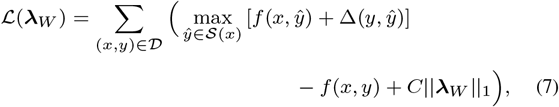

where ‖·‖_1_ is the *ℓ*_1_ norm and *C* is a weight for the *ℓ*_1_ regularization term to avoid overfitting to training data (we used *C* = 0.001 by default). Here, Δ(*y*, *ŷ*) is a loss function of *ŷ* for *y* defined as

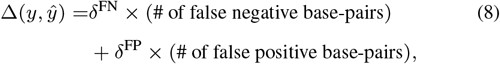

where *δ*^FN^ and *δ*^FP^ are tunable hyperparameters to control the trade-off between sensitivity and specificity for learning the parameters. We used *δ*^fn^ = 8.0 and *δ*^FP^ = 1.0 by default. In this case, we can calculate the first term of Eq. (7) using the Zuker-style dynamic programming algorithm modified by the loss-augmented inference Tsochantaridis *et al.* (2005).

To minimize the objective function (7), we can apply stochastic subgradient descent (Fig. 2) or its variant.

**Fig. 2.**
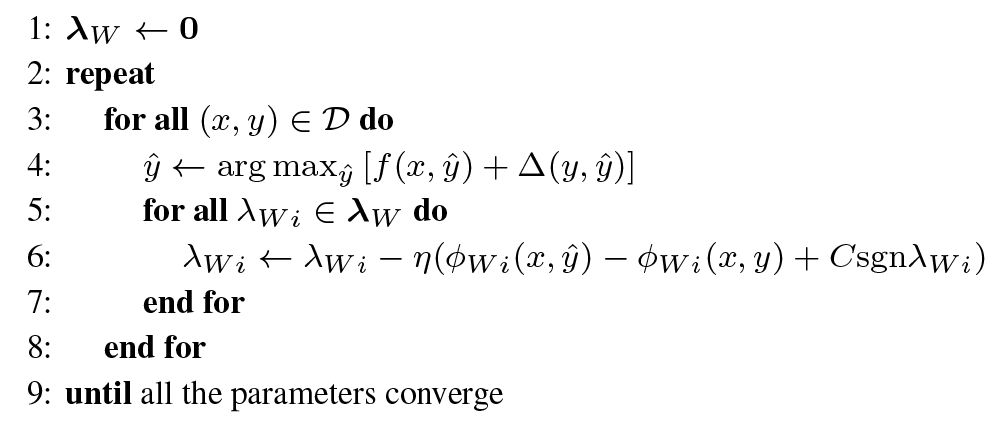
The stochastic subgradient descent algorithm for SSVMs. sgn is the sign function. *η* > 0 is the predefined learning rate.

## 3 Results

### 3.1 Implementation

Our algorithm was implemented as a program called MXfold, which is short for the MaX-margin based rna FOLDing algorithm. The source code is available at https://github.com/keio-bioinformatics/mxfold. The free energy parameters λ_*T*_ was implemented using the Vienna RNA package version 2.3.5 Lorenz *et al.* (2011).

### 3.2 Datasets

In order to evaluate our algorithm, we performed computational experiments on the four datasets assembled by Rivas *et al.* Rivas *et al.* (2012), TrainSetA/TestSetA and TrainSetB/TestSetB. TrainSetA and TestSetA were collected from the literature Dowell and Eddy (2004); Do *et al.* (2006); Andronescu *et al.* (2007); Lu *et al.* (2009); Andronescu*et al.* (2010). TrainSetB and TestSetB were extracted from Rfam Gardner *et al.* (2011), which contain 22 families with 3D structures. The literature-based sets “A” and the Rfam-based sets “B” are structurally diverse. Furthermore, highly identical sequences were removed from all the four datasets. We excluded a number of sequences that contain pseudoknotted secondary structures in the original data sources from all the four datasets since all algorithms evaluated in this study were designed for RNA secondary structure prediction without pseudoknots. The dataset is also available at https://github.com/keio-bioinformatics/mxfold.

### 3.3 Evaluation measures

We evaluated the accuracy of predicting RNA secondary structures through the sensitivity (SEN) and the positive predictive value (PPV), defined as:

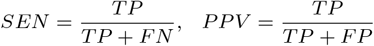

where *TP* is the number of correctly predicted base-pairs (true positives), *FP* is the number of incorrectly predicted base-pairs (false positives), and *FN* is the number of base-pairs in the true structure that were not predicted (false negatives). We also used the F-value as the balanced measure between SEN and PPV, which is defined as their harmonic mean:

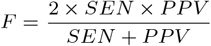

### 3.4 Effects of scoring models

In order to confirm the effects of integration of the thermodynamic model and the machine leaning based model, we performed computational experiments on the datasets described in Sec. 3.2. The trainable parameters of the machine learning based model were trained from TrainSetA. Each model was evaluated with the prediction accuracy of the Viterbi decoding on TestSetA and TestSetB. Table 1 shows the prediction accuracy of three models: the thermodynamic model (TM) that employs only *f*_*T*_ (*x*, *y*) in Eq. (2), the machine learning model (ML) only with *f*_*W*_(*x*, *y*), and our model that integrates the thermodynamic model and the machine learning based model (TM+ML), indicating that our model (TM+ML) performed the most accurate prediction. On TestSetA, our models was slightly better than ML only model. On TestSetB that contains structurally dissimilar RNAs from TrainSetA, the difference of the accuracy between TM+ML and ML is larger.

**Table 1.** The accuracy of each scoring model.

| Model | TestSetA |  |  | TestSetB |  |  |
| --- | --- | --- | --- | --- | --- | --- |
|  | SEN | PPV | F | SEN | PPV | F |
| TM | 0.682 | 0.659 | 0.670 | 0.598 | 0.485 | 0.536 |
| ML | 0.703 | **0.764** | 0.732 | 0.575 | 0.550 | 0.563 |
| TM+ML | **0.715** | 0.761 | **0.737** | **0.617** | **0.565** | **0.590** |

### 3.5 Effects of feature representations

We evaluated the prediction accuracy of the Viterbi decoding on TestSetA and TestSetB for several feature representations. Figure 3 shows the accuracy for each feature representation with different context lengths *L* = {0, 3, 5, 7, 10, 15, 20}. This indicates that the difference of the accuracy on *L* ≥ 7 is negligible although longer sequential features enable more accurate prediction. In addition, as shown in Fig. 4 that shows the running time for each context length, sequential features of longer context lengths need more calculation time. Therefore, we set the default context length *L* = 7 since shorter sequential features decrease the number of trainable features reducing the risk of overfitting.

**Fig. 3.**
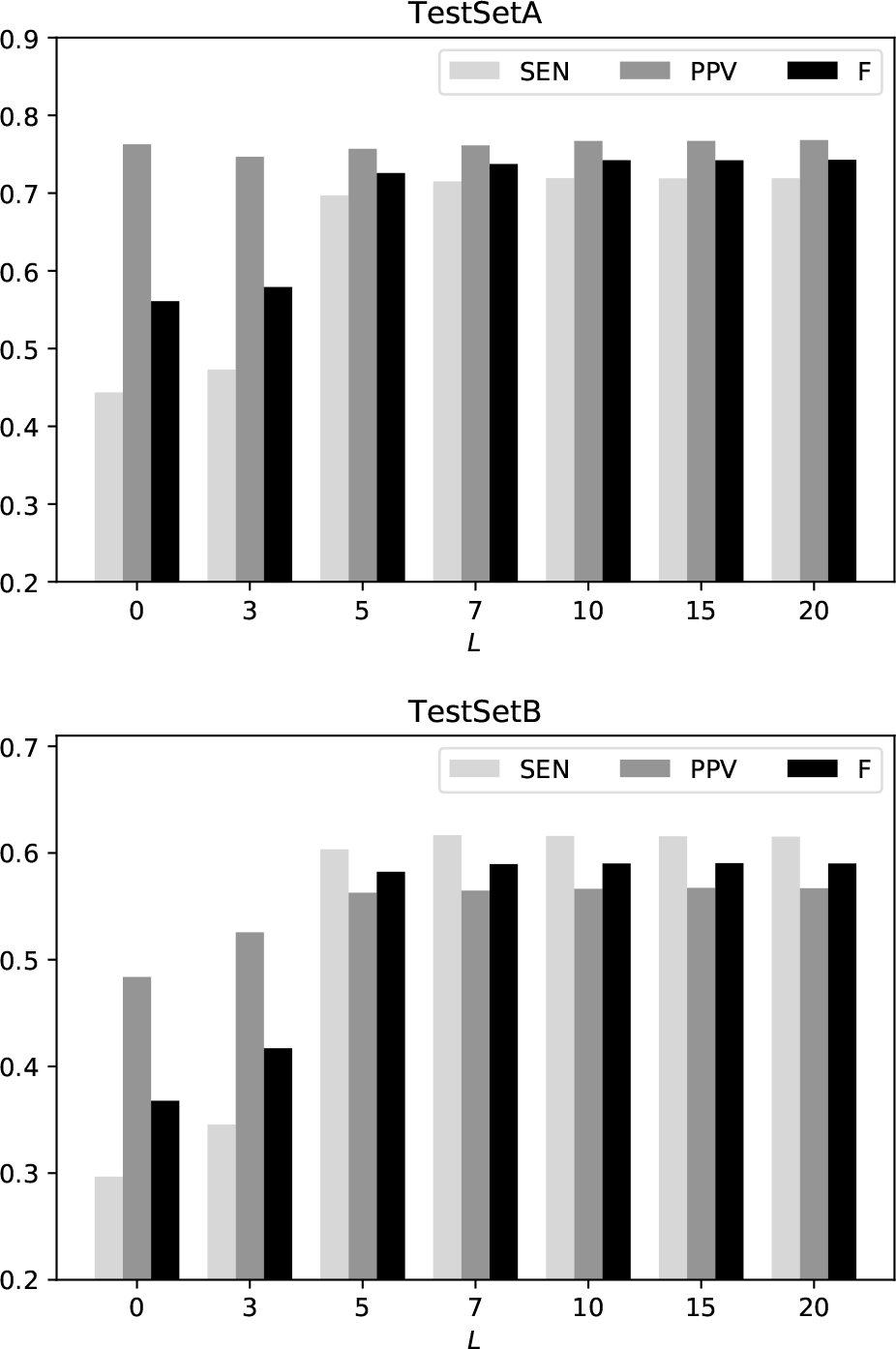
The accuracy for each feature representation with different context lengths *L* on TestSetA (top) and TestSetB (bottom).

**Fig. 4.**
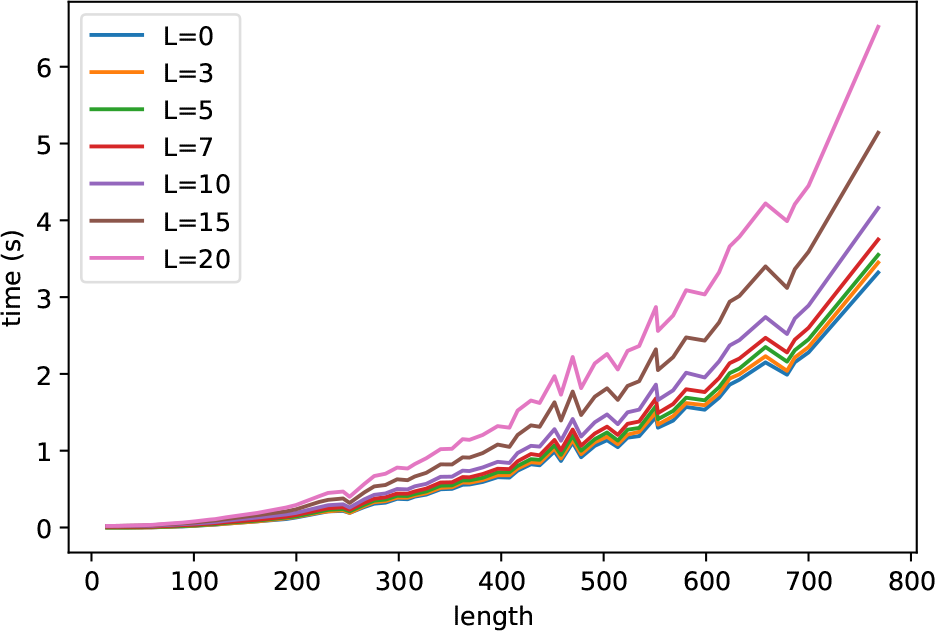
The running time for each feature representation with different context lengths *L* measured on Linux OS v2.6.32 with Intel Xeon E5-2680 (2.80 GHz) and 64 GB memory.

### 3.6 Comparison with competitive methods

We compared our algorithm with the competitive methods including CentroidFold version 0.0.15 Hamada *et al.* (2009); Sato *et al.* (2009), CONTRAfold version 2.02 Do *et al.* (2006), RNAfold in the Vienna RNA package version 2.3.5 Lorenz *et al.* (2011) and ContextFold version 1.00 Zakov *et al.* (2011). For the posterior decoding methods with the trade-off parameter *γ* in Eq. (5), we used *γ* ∈ {2^*n*^ | *n* ∈ ℤ, −5 ≤ *n* ≤ 10}.

Figure 5 shows PPV-SEN plots for each method, indicating that our algorithm works accurately on TestSetA and TestSetB. On TestSetA, ContextFold (F=0.742) is slightly better than MXfold with Viterbi decoding trained from TrainSetA (F=0.737). Whereas, on TestSetB, ContextFold (F=0.496) is much worse than MXfold with Viterbi decoding trained from TrainSetA (F=0.590) and others. Furthermore, MXfold with Viterbi decoding trained from both training datasets performed the most accurate prediction (F=0.626).

**Fig. 5.**
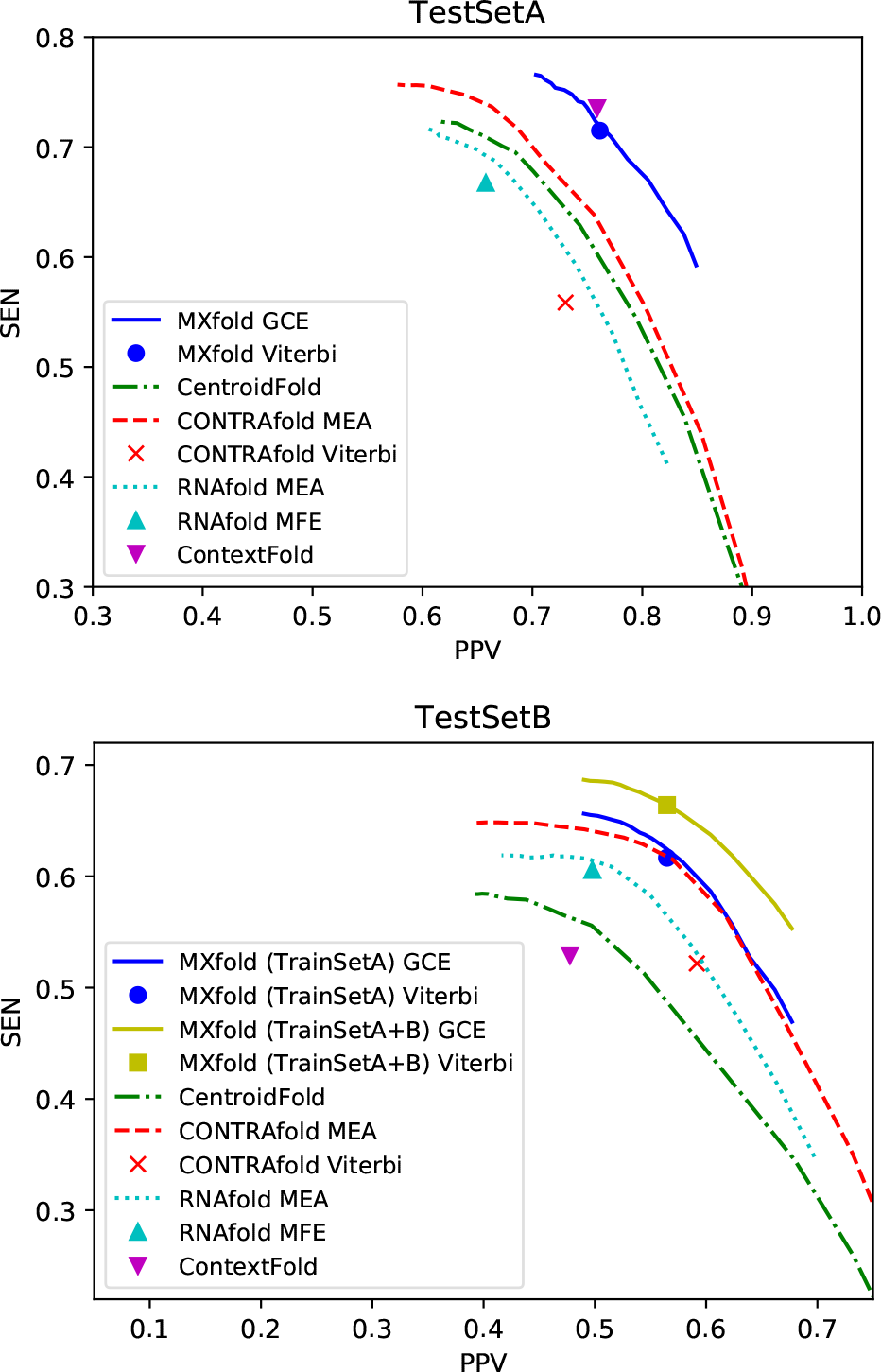
PPV-SEN plots comparing our algorithm with the competitive methods on TestSetA (top) and TestSetB (bottom).

Figure 6 shows the running time for each method for the lengths of input sequences in TestSetA, indicating that our algorithm with the Viterbi decoding is comparable with the other methods in the running time although our algorithm with the posterior decoding is much slower than the other methods.

**Fig. 6.**
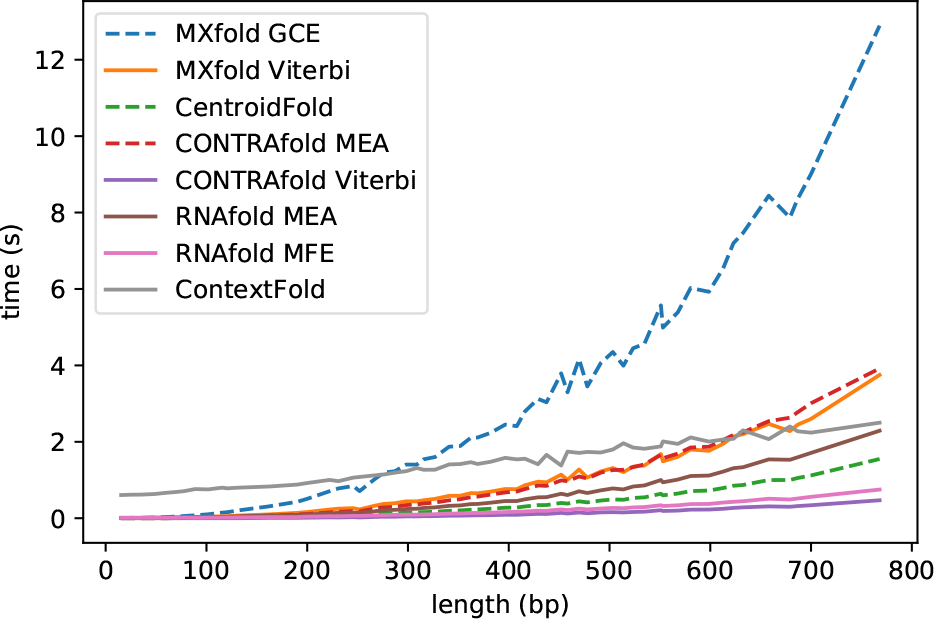
The running time for the lengths of input sequences measured on Linux OS v2.6.32 with Intel Xeon E5-2680 (2.80 GHz) and 64 GB memory.

## 4 Discussion

Table 1 compares the three models: the thermodynamic model (TM), the machine learning based model (ML) and the integrated model (TM+ML). Since the thermodynamic model *f*_*T*_(*x*, *y*) is implemented using the Vienna RNA package, the prediction result of TM is similar to that of RNAfold. The result on TestSetA indicates that the difference between ML and TM+ML is very small. We can explain that this is because the trainable parameters of ML and TM+ML are identical to each other, and the learning algorithm works well on both models. On the other hand, since the literature-based TrainSetA and the Rfam-based TestSetB are structurally diverse as described in Sec. 3.2, TestSetB includes a number of substructures whose scoring parameters cannot be trained from TrainSetA. TM+ML model can calculate scores for such “unobserved” substructures using the thermodynamic energy parameters although ML only model cannot. Our integrated model can improve the prediction accuracy by complementing missing parts each other.

We compared the learnability of our model for several context lengths *L* of sequential features in Fig. 3. Most existing models including RNAfold and CONTRAfold use the context length 3 ≤ *L* ≤ 5, whose accuracy shown in Fig. 5 is close to that of our model with the same range of the context length. Although Fig. 3 shows that longer context length of sequential features enables us to improve the prediction accuracy, its effects tend to be saturated at *L* = 7. The objective function of our algorithm (7) contains the *ℓ*_1_ regularization term, by which rarely used parameters (e.g., sequential features with *L* > 7) quickly shrink toward zero at line 6 of Fig. 2. Hereby, the risk of overfitting caused by rarely observed features can be reduced.

Figure 5 shows that ContextFold achieved the best accuracy on TestSetA, but the worst on TestSetB. Similarly, the accuracy of CentroidFold on TestSetB remarkably deteriorated compared with that on TestSetA. The common point between ContextFold and CentroidFold is the training data: ContextFold and the Boltzmann likelihood (BL) parameter set used in CentroidFold were trained from the S-Full dataset Andronescu *et al.* (2010), which is one of the datasets included in TrainSetA. This suggests that ContextFold and the BL parameter set fell into the overfitting. There is a possibility that ContextFold trained from TrainSetA+B achieves more accurate prediction than MXfold trained from TrainSetA+B. However, ContextFold might not work well for other sequences dissimilar from TrainSet A and B because of the overfitting. Meanwhile, we can expect that our algorithm that integrates the thermodynamic model still performs robust and accurate prediction without overfitting for such sequences due to the integrated thermodynamic model.

The posterior decoding algorithms are known to be one of effective approaches for many combinatorial optimization problems Carvalho and Lawrence (2008). In fact, the posterior decoding with CONTRAfold (MEA) achieves much better accuracy than its counterpart of the Viterbi decoding as shown in Fig. 5. However, we can surprisingly observe no advantage for the posterior decoding for MXfold (GCE). CONTRAfold was trained by the conditional log-linear models (CLLMs) in which the expectation of the occurrence of features is used for calculating gradients of the objective function. The posterior decoding algorithms employ the base-pairing probabilities that are also calculated by the expectation of the occurrence of base-pairs. This can be interpreted that the optimization with CLLMs is appropriate for the posterior decoding. SSVM used by our algorithm considers only the optimal structure with the (loss augmented) Viterbi algorithm for each training step. This means that SSVM is optimized for the Viterbi decoding, but not for the posterior decoding that considers not only the optimal structures but also the distribution of all possible structures. As shown in Fig. 6, the posterior decoding algorithms are much time-consuming compared with their counterparts of the Viterbi and MFE algorithms. Therefore, although the posterior decoding with the parameters learned by CLLMs is one of the best solution from the viewpoint in the prediction accuracy, the Viterbi algorithm with SSVM is a practical alternative.

## 5 Conclusion

We proposed a novel algorithm for RNA secondary structure prediction that integrates the thermodynamic approach and the machine learning based weighted approach. Our fine-grained model combines the experimentally determined thermodynamic parameters with a large number of scoring parameters for detailed contexts of features that are trained by the structured support vector machine (SSVM) with the *ℓ*_1_ regularization to avoid overfitting. Our benchmark shows that our algorithm achieves the best prediction accuracy compared with existing tools, and heavy overfitting as shown in ContextFold cannot be observed.

Accurate secondary structure prediction for long RNA sequences has been demanded since long non-coding RNAs (lncRNAs) have recently been emerging. To respond to such demand, we need to implement the sparsification technique Backofen *et al.* (2011) to our algorithm with the Viterbi decoding. As shown in Fig. 6, ContextFold that implements the sparsification technique enables us fast structure prediction even for long sequences.

The base-pairing probabilities calculated from the posterior distribution have been required for various applications for RNA informatics such as family classification Sato *et al.* (2008); Morita *et al.* (2009), pseudoknotted RNA secondary structure prediction Sato *et al.* (2011), RNA-RNA interaction prediction Kato *et al.* (2010) and simultaneous aligning and folding Sato *et al.* (2012). Accurate base-pairing probabilities calculated by our algorithm can improve the quality of such applications.

## Acknowledgements

The supercomputer system was provided by the National Institute of Genetics (NIG), Research Organization of Information and Systems (ROIS).

*Conflict of Interest:* None declared.

